# CD8 co-receptor enhances T cell activation without any effect on initial attachment

**DOI:** 10.1101/2020.05.05.079145

**Authors:** Philippe Robert, Laurent Limozin, Anton van der Merwe, Pierre Bongrand

## Abstract

The scanning of surrounding tissues by T lymphocytes to detect cognate antigen is a highly demanding process that requires high rapidity, sensitivity and specificity. Co-receptors such as CD8 are known to increase detection performance, but the exact mechanism of this role remains incompletely understood. Here, we used interference reflection microscopy to image the initial spreading of 1G4 receptor transfected CD8+ and CD8− Jurkat cells dropped on surfaces exposing five cognate antigens of varying activating power, and we used a laminar flow chamber to measure the influence of CD8 on the kinetics of bond formation and rupture between cell-born T cell receptors (TCRs) and peptide-exposing major histocompatibility complex antigens (pMHCs) at the single molecule level. It is concluded that CD8 did not influence TCR-pMHC interaction during the first seconds following cell surface encounter, but it promoted the spreading responses during the first minutes, thus suggesting that CD8 was involved in early activation rather than binding. In addition, presented results were quantitatively compared with a recent report on the cell-free interaction between the same ligand-receptor couples : it is concluded that bond formation was strongly impaired by cell molecular environment, while bond rupture was comparable in both systems. Results from this and previous reports were used to propose a quantitative scheme of the strategy used by T lymphocytes to scan foreign surfaces. It is suggested that the understanding of the strategy used by cells to perform their basic functions may be a prerequisite to understand the function of molecular networks revealed by high throughput methods.

## Introduction

The detection by T lymphocytes of foreign peptides (p) bound to major histocompatibility complex (MHC)-encoded proteins (pMHC) is a key and early step of immune responses. T cell receptors (TCRs) are highly specific since they can detect a cognate peptide surrounded by a large excess of molecules differing by a few or even a single amino acid (*van der Merwe 2011*, *Malissen 2015*). Detection is very sensitive since it can be triggered by a few or even a single pMHC. It is also very rapid since signaling events such as tyrosine phosphorylation, a rise in cytoplasmic calcium concentration, diacylglycerol production, or retraction of TCR-rich microvilli may occur within a few seconds (*Huse 2007*, *Brodovitch 2013*, *Cai 2017*). Last but not least, this initial event can generate a wide range of cellular outcomes including expression of activation antigens such as CD69, production of mediators such as IL-2, triggering of differentiation or proliferation programs, rapid destruction of a target cell, or even T cell inactivation. It would be of obvious theoretical and practical interest to gain an accurate understanding of the links between molecular events involved in this process. While an impressive amount of information has been obtained in the last few years, our understanding of the activation process remains incomplete and some key information is lacking.

Much experimental evidence supports the hypothesis that the discrimination process is dependent on the physical properties of TCR-pMHC interaction : it was first reported that activation generally required that the life-time of this interaction be higher than a few seconds when assayed with soluble molecules, e.g. by using surface plasmon resonance (*Matsui 1994*). However, some discrepancies remained and it was recognized that so-called 3D interactions involving soluble molecules were not representative of phenomena occurring on the cell membrane, since under physiological conditions bonds might be subjected to mechanical forces likely to modulate their lifetime. Definite progress was obtained when the TCR-pMHC interaction was studied a the **single molecule level** with methods based on laminar flow chambers (*Robert 2012*, *Limozin 2019*) or micropipette assays (*Liu 2014*). Experimental results suggested that the activation potency of a TCR-pMHC interaction was closely linked to its capacity to resist a pulling force on the order of 10 pN, which could be generated by the cell machinery (*Liu 2014*). Another important point is that T cells were suggested to integrate short binding events during a period of time ranging between 30s and perhaps more than 1,000s depending on the studied signal (*Brodovitch 2013*, *Liu 2014*, *Lin 2019*). As a consequence, the rate of bond formation should be considered as well as bond lifetime.

Another limitation of aforementioned experiments is that they did not address the potential role of co-stimulatory molecules in the triggering process. Indeed, the sensitivity of antigen detection is strongly enhanced by co-receptors such as CD8, and this may involve many non exclusive mechanisms such as enhancement of intercellular adhesion, facilitating TCR-pMHC interaction, altering the conformation of the trimolecular complex CD8-pMHC-TCR, or promoting subsequent signaling events, e.g. by recruitment of CD8-interacting kinase p56lck (*Smith-Garvin 2009*). Indeed, a molecule such as CD8 has been shown to induce adhesion between CD8 T cells and cells expressing sufficient amounts of class I MHC (*Norment 1988*). In vitro studies suggested that CD8 could enhance the rate of bond formation between TCR and pMHC without influencing bond lifetime (*Dutoit 2003*, *Gakamsky 2005*). In aforementioned reports on correlation between T cell activation and lifetime of TCR-pMHC bonds, Liu et al. used a mutated MHC to prevent CD8 binding (*Liu 2014*) and Robert et al. (*Robert 2012*, *Limozin 2019*) used a cell-free system including only TCR and pMHC. However, other experiments suggested that CD8 might significantly increase the resistance of TCR-pMHC interaction by forming a trimolecular complex (*Jiang 2011*, *Kolawole 2018*, *Hong 2018*), but this occurred after a lag of 1 second or more (*Jiang 2011*, *Hong 2018*) and required tyrosine phosphorylation events. Since phosphorylation is a well known key step of T lymphocyte activation (*Smith-Garvin 2009*), and this may involve the p56lck Src kinase associated with CD8, it is not clear whether CD8 might enhance lymphocyte activation by potentiating signaling events with a consecutive binding enhancement, or it might enhance signaling events independently of the binding process.

Here, we investigated the potential role of CD8 in modulating TCR-pMHC interaction and triggering early activation events in a cellular model. We compared the interaction of CD8− and CD8+ Jurkat cells transfected with 1G4 TCR and surfaces coated with pMHC complexes of graded activation potency as revealed by measuring interferon production (*Aleksic 2010*). The effect of CD8 on TCR-pMHC molecular interaction was assayed with a laminar flow chamber, allowing us to explore the efficiency of bond formation during encounters of a few millisecond duration, and bond lifetime during a monitoring period of several seconds (*Robert 2012*, *Limozin 2019*). As a cell response, we measured the rate and extent of spreading during the first 15 minutes following contact with antigens. Indeed, it was previously found that the maximum spreading rate determined during the first 3 minutes after contact was robustly correlated to pMHC activation potency (*Brodovitch 2015*).

## Materials and Methods

### Cells

As previously, we used a human Jurkat T cell line expressing the 1G4 TCR (*Brodovitch 2015*). As this line does not express CD8 (CD8−), we used lentiviral transduction (*Denham 2019*) to produce a version that expresses CD8αβ (CD8+).

### Molecules and surfaces

The wild type or H74A mutant of HLA-A2 heavy chain (residues 1-278) with C-terminal BirA tag and β2-microglobulin were expressed as inclusion bodies in E.coli, refolded in vitro in the presence of synthesized variants of the NY-ESO-1_157–165_ peptide SLLMWITQV (9V), and purified using size-exclusion chromatography (*Chen 2005*, *Aleksic 2010*). The pMHCs used in this study, as in previously reported experiments (*Brodovitch 2015*, *Limozin 2019*), were 9V and variants 3A, 3Y, or 9L in complex with wild-type HLA-A2, or 9V in complex with the HLA-A2 mutant H74A.

Glass surfaces were prepared as previously described (*Brodovitch 2015*, *Limozin 2019*) by sequential cleaning with a mix of 70% sulfuric acid and 30% H_2_0_2_, coating with poly-L lysine (Sigma-Aldrich, 150,000-300,000 MW), activation with glutaraldehyde (2.5%), coupling with biotinylated-bovine serum albumin (100 μg/ml, Sigma) then neutravidin (10μg/ml), and finally coating with biotinylated pMHCs. Absolute calibration of pMHC surface density was done as previously described by labeling with an excess of Alexa Fluor 488 labeled anti-HLA antibody (Biolegend, San Diego) and fluorescence determination.

### Cell spreading experiments

Experimental procedure was previously described (*Pierres 2003*, *Pierres 2008*, *Brodovitch 2015*). Briefly, 0.5 ml of cell suspension in HEPES-buffered RPMI medium suspended with 10% fetal calf serum were deposited in teflon-walled wells maintained at 37°C on the stage of an inverted microscope (Axiovert 135, Zeiss) bearing a heating enclosure (TRZ 3700, Zeiss) set at 37°C. Interference reflection microscopy (IRM), also called Reflection interference contrast microscopy (RICM) was performed with a 63x Antiflex objective, 546 nm excitation light and an Orca C4742-95-10 camera (Hamamatsu, Japan). Pixel size was 125×125 nm^2^. In a typical experiment, the microscope was set on a random field and about 600 images were recorded with 1Hz frequency. Thus the initial contact and spreading of between 5 and 10 cells could be monitored (Figure 1). Image stacks were processed with a custom-made software (*Pierres 2008*) that performed mean filtering (25-pixel areas, for noise reduction), linear compensation for inhomogeneities of field illumination and temporal variations of light intensity. Cell/substratum distance d at each pixel was derived from illumination intensity I with the low incidence approximation :

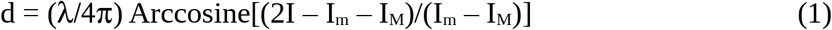

where λ is the light wavelength, and I_m_ and I_M_ are, respectively, the minimum and maximum intensities corresponding to d=0 and d = λ/4 ~ 126 nm, respectively. These intensities were determined by assuming that all distances occurred somewhere during the 10 min observation period. Molecular contact between cells and surfaces was assumed to occur when the calculated distance d was less than 34 nm, based on the added size of interacting receptors. It was checked that dark zone were indicative of *bona fide* attachment by assessing cell stability in presence of a low hydrodynamic flow (*Pierres 2003*).

**Figure 1 :**
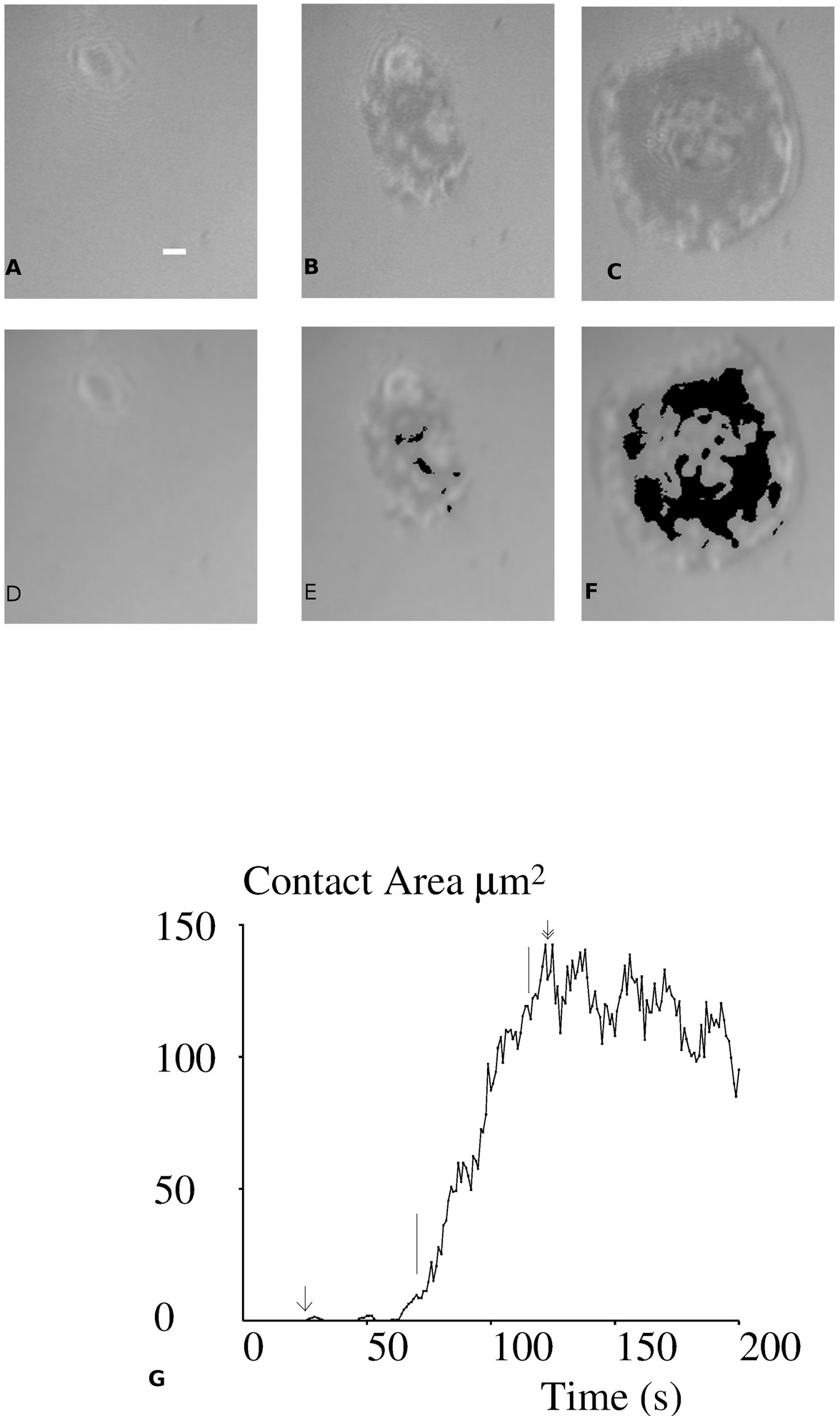
spreading of a cell on an activating surface. Jurkat cells expressing 1G4 TCR were dropped on a surface coated with 3Y pMHC and a random field was monitored with IRM/RICM. Images of a typical cell are shown at time 0 (A), 64 s (B) and 100 s (C). The contact areas are shown in black (D, E, F) : they were estimated at 0 (D), 2.8 (E) and 87 μm^2^. (F) The spreading curve is also shown : maximal spreading area is 142 μm^2^, the maximal spreading rate is 2.7 μm^2^/s. Bar is 2 μm length..

When the 10-minute recording was completed, Images of typically 10 random fields were rapidly recorded in order to obtain better statistics for final contact area. A total of 396 spreading kinetics and 11,161 final spreading areas were recorded on 1G4C-D8+ cells. They were obtained under the same conditions as the 495 spreading kinetics and 13,574 spreading areas previously obtained on 1G4-CD8− cells.

Spreading kinetics were analysed with a custom-made software as follows : time zero was defined as the first time where contact was larger than or equal to 2 pixels on corrected images, corresponding to an area of 0.03 μm^2^. The slope of area versus time curve was then calculated on a sliding window of 45 second duration, which yielded the maximum spreading rate. The lag before spreading was defined as the difference between the onset of maximum spreading rate and time zero. Finally, the maximum contact area was determined over the 10 minute monitoring time. A typical spreading curve is displayed on Figure 1.

### Cell adhesion under flow

We used a previously described laminar flow chamber setup (*Robert 2011*) and data analysis was performed with an improved software that was recently applied on microspheres (*Limozin 2019*). Here, cells were driven along surfaces coated with various pMHC densities by a laminar flow. Wall shear rate was U=16.6 ± 4.4 s^−1^ (SD, n=38 experiments), and the average velocity U of sedimented cells (this is the so-called hydrodynamice velocity) was 62 μm/s. The motion of cells crossing a microscope field was recorded with 25 Hz frequency. A cell was considered as arrested when the centroid position moved by less than 1.6 μm (i.e. 2 pixels) during a 0.32 s time interval.

Assuming that arrests were mediated by the interaction of single microvilli with surface-bound pMHCs, the force F on the bond was calculated by modeling cells as spheres, yielding (*Pierres 1995*, *Limozin 2019*)

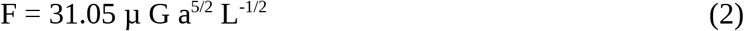

where μ is the medium viscosity (0.91 10^−3^ Pa.s at 25°C), a is the cell radius (6.8 μm) and L is the microvillus length (estimated at about 1 μm, note that F is only weakly dependent on the precise value of L). F was thus estimated at about 55 pN. It may be noticed that, due to a torque effect, the force on the bond is higher than the force on the cell that is about 22 pN.

Another result of fluid mechanics (*Goldman 1967*), the relevance of which to cells was subjected to experimental check (*Tissot 1992*), is that the relative velocity of the sphere and chamber surfaces near contact is on the order of 0.43U ≈ 7.1 μm/s. This yields a maximum value of 8 ms for the duration of TCR-pMHC interaction if these molecules are modeled as freely moving rods of 15 nm length. It must be emphasized that this is only an upper bound for the actual encounter time, since this is dependent on the distance between surfaces.

In the present work, cell motion was followed along a total path of about 13.6 meters, and a total number of 3,936 arrests were recorded. Arrests were monitored for at least 5 seconds, and results were used to build survival curves, yielding a quantitative assessment of bond lifetime.

The binding linear frequency (BLD) was defined as the number of detected cell arrests per unit of distance traveled at the velocity corresponding to sedimented cells.

### Statistics

Statistical calculations were performed following standard procedures (*Snedecor & Cochran, 1980*). Briefly :

Since cell arrests in the flow chamber appeared as rare events with a frequency lower than 1 per 1,000 pixels (800 μm), they were assumed to follow Poisson statistics, and the relative uncertainty of arrest frequency was estimated as 1/N^1/2^ in an experiment amounting to a total number of N arrests.

Since the main purpose of this work was to compare the behavior of two cell populations (1G4CD8− and 1G4-CD8+) under a number of conditions (five pMHC, 3 to 4 surface densities), the non-parametric signed-rank test was first used before performing more quantitative comparisons. It must be emphasized that this test does not require that the distribution of tested parameters be normal, and its sensitivity was claimed to be at least higher than 86% of t-test sensitivity and sometimes higher (*Snedecor & Cochran*, p 146)

The effect of CD8 on bond lifetime was analyzed with Chi2 tests by comparing the frequencies of arrest durations falling in five time intervals ([in seconds : [0,0.3[, [0.3,0.75[, [0.75,1.5[, [1.5,5[, [5,∞[).

Finally, quantitative parameters such as contact areas or spreading rates were compared with Student’s t-test, using Satterthwaite’s correction for unequal samples. Standard Pearson correlation coefficients were also calculated to assess correlations between different parameters.

## Results

### Spreading experiments

#### Cells display a typical spreading pattern

In a series of experiments, 1G4-CD8+ cells were dropped on surfaces coated with four different surface densities of the five pMHC species, amounting to 20 conditions. In each experiment, a microscope field was randomly selected and monitored with IRM/RICM for 10 minutes. Images were recorded with 1 Hz frequency. A typical example is shown on Figure 1 : initial cell-surface encounter was defined as the first occurrence of a 2-pixel (1/32 μm^2^) contact area. Several tens of seconds later, the contact area started increasing rapidly. This process was quantified by determining the 45s period of time with maximum area increase : the average slope of the time/area plot (Figure 1G) during this time interval was defined as the maximum spreading rate (in μm^2^/s) and the spreading lag was defined as the time difference between the onset of maximum spreading velocity and initial contact. Finally, the contact area reached a maximum after several tens of seconds, and displayed very slow decrease during the following ten minutes. When the ten minute observation period was completed, a microscope image in transmitted light was recorded, and at least 10 supplementary random fields were selected for recording of both visible and IRM/RICM images (Figure 2). This allowed more than tenfold increase of the number of monitored cells. Indeed, a total number of 309 spreading series could be analyzed, and 13,574 additional static cell images were obtained. The latter series were used to calculate the spreading area and fraction of spreading cells (Figure 2).

**Figure 2 :**
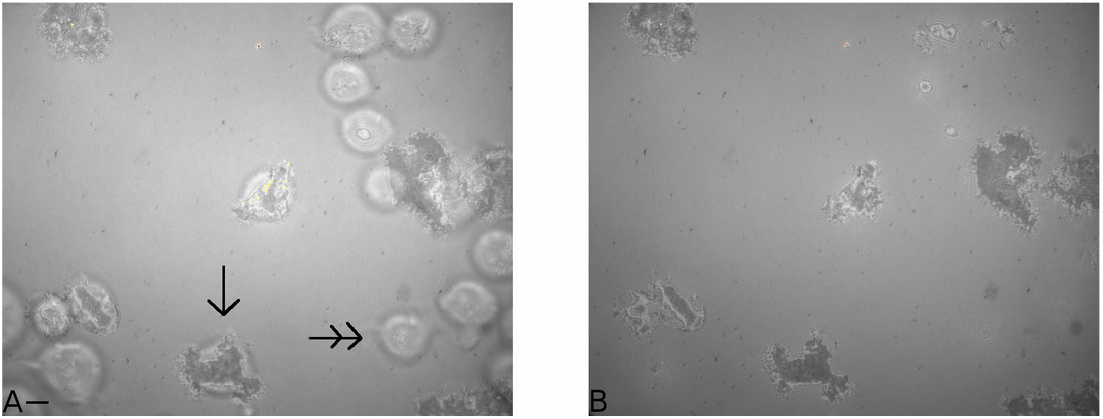
Only a fraction of cells display spreading behaviour. A typical microscope field is shown (time : 600 second, 3Y-coated surfaces). A : Conventional and IRM/RICM images are superimposed. B : IRM/RICM image. There is a clearcut difference between spread cells (arrow) and cells that did not form noticeable contact (double arrow). Bar is 5 μm length.

First, we studied the correlations between measured parameters. As shown on Figure 3, the maximum spreading area and the maximum spreading rate were strongly correlated (Figure 3A, P< 10^−7^). However, the lag before spreading and maximum spreading rate were only weakly and negatively correlated (Figure 3B, P≈0.02). In addition, quantitative analysis of the static images recorded under 20 conditions (5 peptides and 4 surface concentrations) led to the conclusion that the spreading area and the fraction of spreading cells were strongly and positively correlated (P<0.01), while they were not correlated to the lag before rapid spreading. Thus, CD8+ cells behaved similarly as CD8− cells (*Brodovitch15*).

**Figure 3 :**
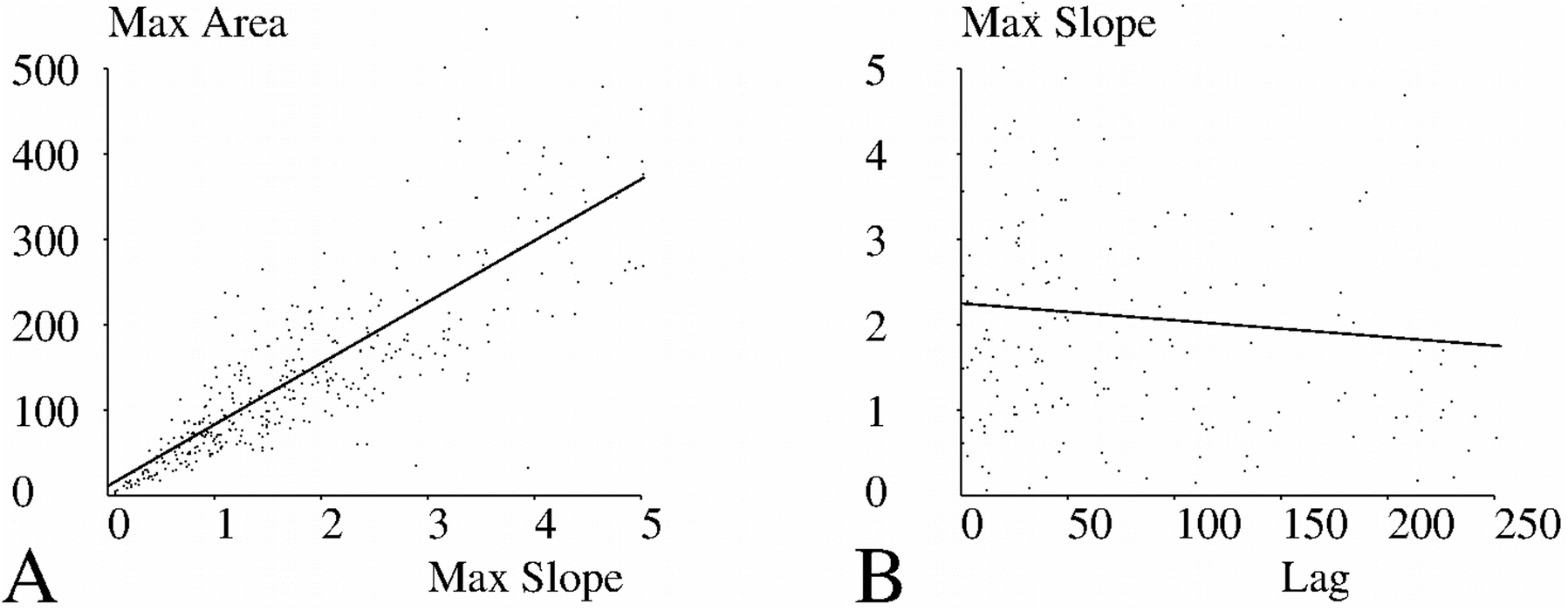
Relationship between spreading parameters. CD8+ 1G4+Jurkat cells were deposited on glass surfaces coated with different amounts of peptide ligands of 1G4. A total of 20 experimental conditions (5 peptides, 4 surface concentrations) were explored A : The individual values of maximum contact area and maximum rate of area increase obtained by processing 379 curves are shown. The correlation coefficient beween displayed parameters was r=.836, in accordance with the strong correlation that is apparent on the figure. B : The individual values of maximum rate of area increase and lag between initial contact and maximum spreading rate obtained by processing 183 curves. The correlation coefficient beween displayed parameters was r=.0952, in accordance with the lack of correlation suggested by the figure

Results obtained with CD8+ and CD8− cells (40 conditions) were then processed to obtain some insight into the spreading process : interestingly, peptide concentration was significantly correlated to spreading area (P=0.02), maximum spreading rate (P=0.005), and fraction of spreading cells (P=0.02), not to the lag between initial contact and spreading initiation (P=0.8), in accordance with the hypothesis that the duration of the process preceding contact was driven by an internal cell mechanism.

#### CD8+ cells display higher spreading fraction, spreading rate and spreading area than CD8− cells

First, we used the sign test to perform a global assumption-free comparison of the spreading behaviour of CD8− and CD8+ cells : CD8+ cells displayed higher maximum spreading rate (P=0.041), higher spreading area (P=0.0004), and higher fraction of cells that spread under similar conditions (P=0.0026). Unexpectedly, CD8+ cells displayed longer delay between contact and onset of rapid spreading (P=0.012).

Then, we asked whether the differences between the spreading behaviour of CD8− and CD8+ cells were dependent on the quality and surface density of activating pMHCs. Since a total of 20 comparisons were performed for each parameter, a difference between CD8+ and CD8− cells was deemed significant only when the significance level was at most 0.0025, i.e. 1/20th of the usually considered 0.05 level.

Firstly, we compared the lag between contact time and onset of rapid spreading : As shown on Figure 4, the only significant finding was that the lag was higher with CD8+ cells than CD8− cells when the activating peptide was H74, and the lowest surface density was used.

**Figure 4.**
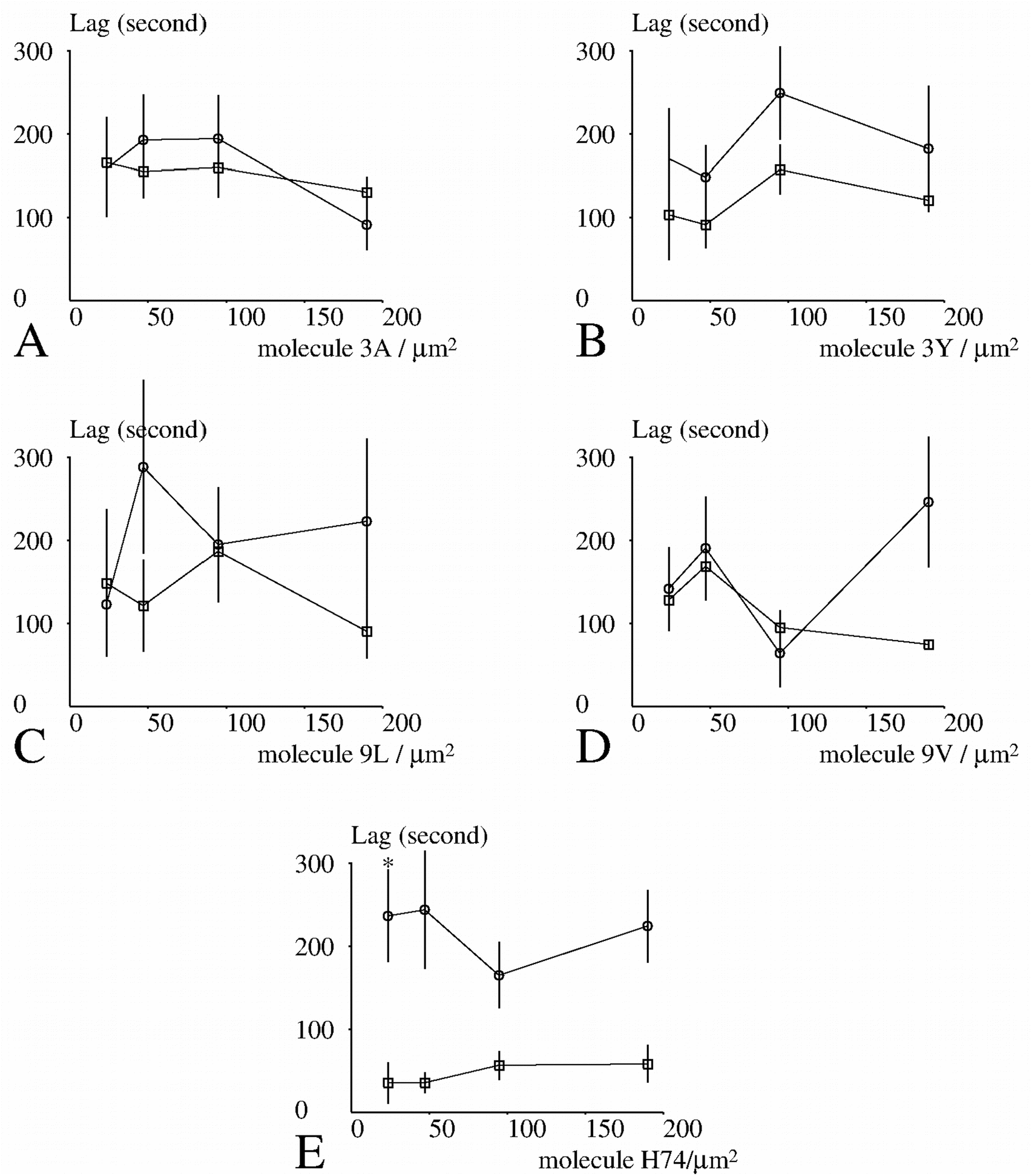
Dependence of the lag between contact and spreading initiation on experimental conditions. CD8+ (open circles) or CD8− (open squares) cells were deposited on surfaces coated with different amounts of TCR ligands. Individual cells were monitored for quantitative determination of the kinetics of contact formation. A total of 549 curves could be processed (which required accurate active spreading and clearcut determination of the time of initial contact formation) and average values of the lag between initial contact and maximum spreading rate are shown. Vertical bar length is twice the standard error for the mean. The significance of differences was calculated with Student’s t test with Satterthwaite correction. Stars indicate a significance P<0.0025.

Secondly, we compared the maximum spreading rates of both cell populations under different conditions : as shown on Figure 5, the only significant difference was found with 3A peptide and the lowest surface density.

**Figure 5.**
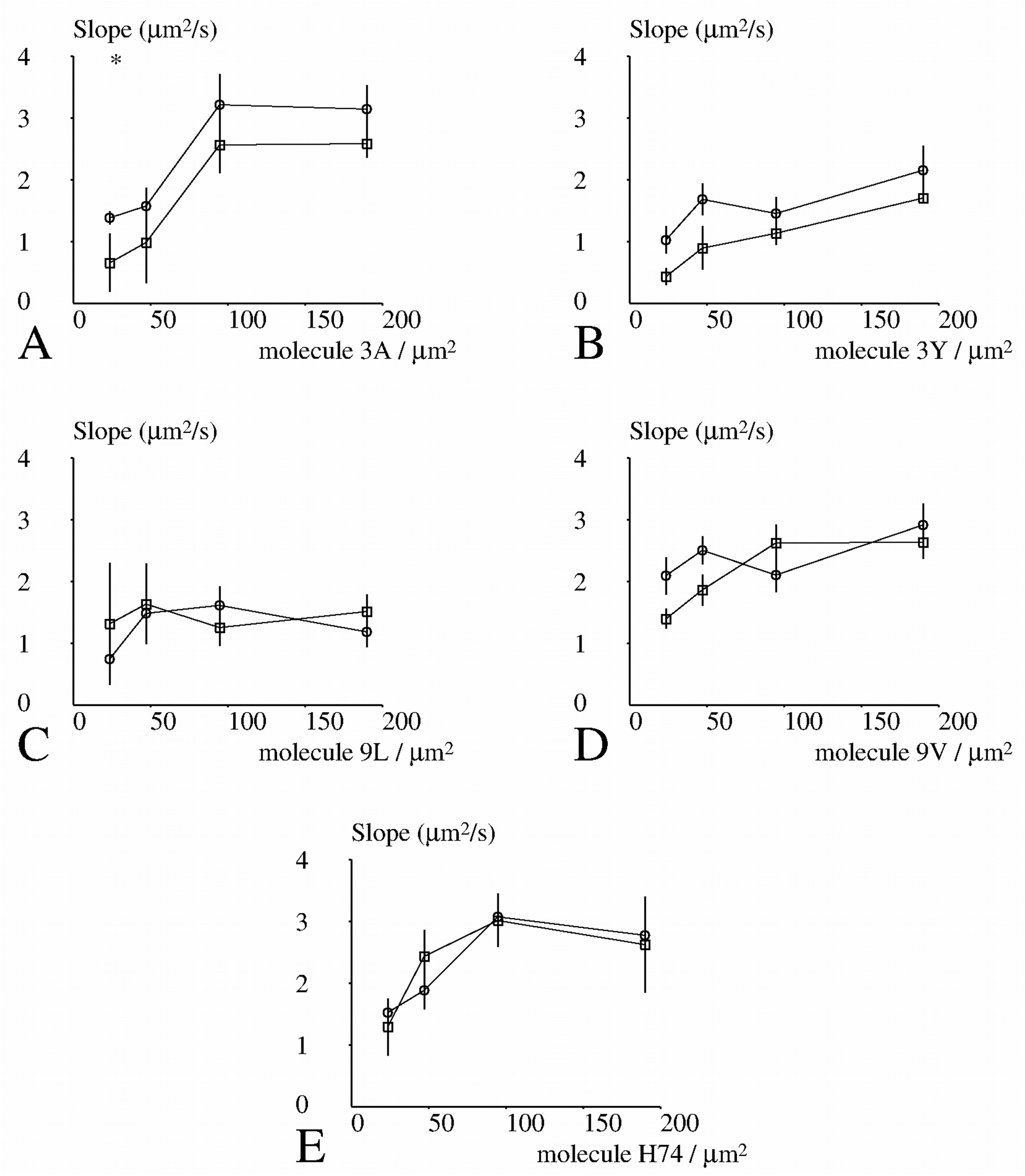
Dependence of the maximum spreading rate on CD8 and activating peptide. CD8+ (open circles) or CD8− (open squares) cells were deposited on surfaces coated with different amounts of TCR ligands. Individual cells were monitored with IRM/RICM for quantitative determination of the kinetics of contact formation. A total of 694 curves could be processed and average values are shown. Vertical bar length is twice the standard error for the mean. The significance of differences was calculated with Student’s t test with Satterthwaite correction. Stars indicate a significance P<0.001.

It must be emphasized that the above two parameters could only be determined with complete spreading kinetics : Since 40 conditions were studied (two cell types, five peptide species and four concentrations), and 891 kinetic plots were obtained, the average number of observations per point was about a dozen, which was not very high. Much more reliable statistics could be obtained with the following two parameters (mean spreading area and fraction of spreading cells), since comparisons were based on 24735 cells, i.e. nearly 600 experimental values per condition.

As shown on Figure 6, CD8+ cells displayed generally higher spreading area than CD8− cells, and results were particularly reproducible for the lowest peptide surface densities. Also, while the least activating peptides 9L and 3Y were reported to trigger only minimal spreading of CD8− cells, whatever the peptide density (*Brodovitch 2015*), CD8+ cells displayed substantial spreading on 3Y, and the spreading area measured on 9L, while fairly low, was about two-fold higher than that of CD8− cells.

**Figure 6.**
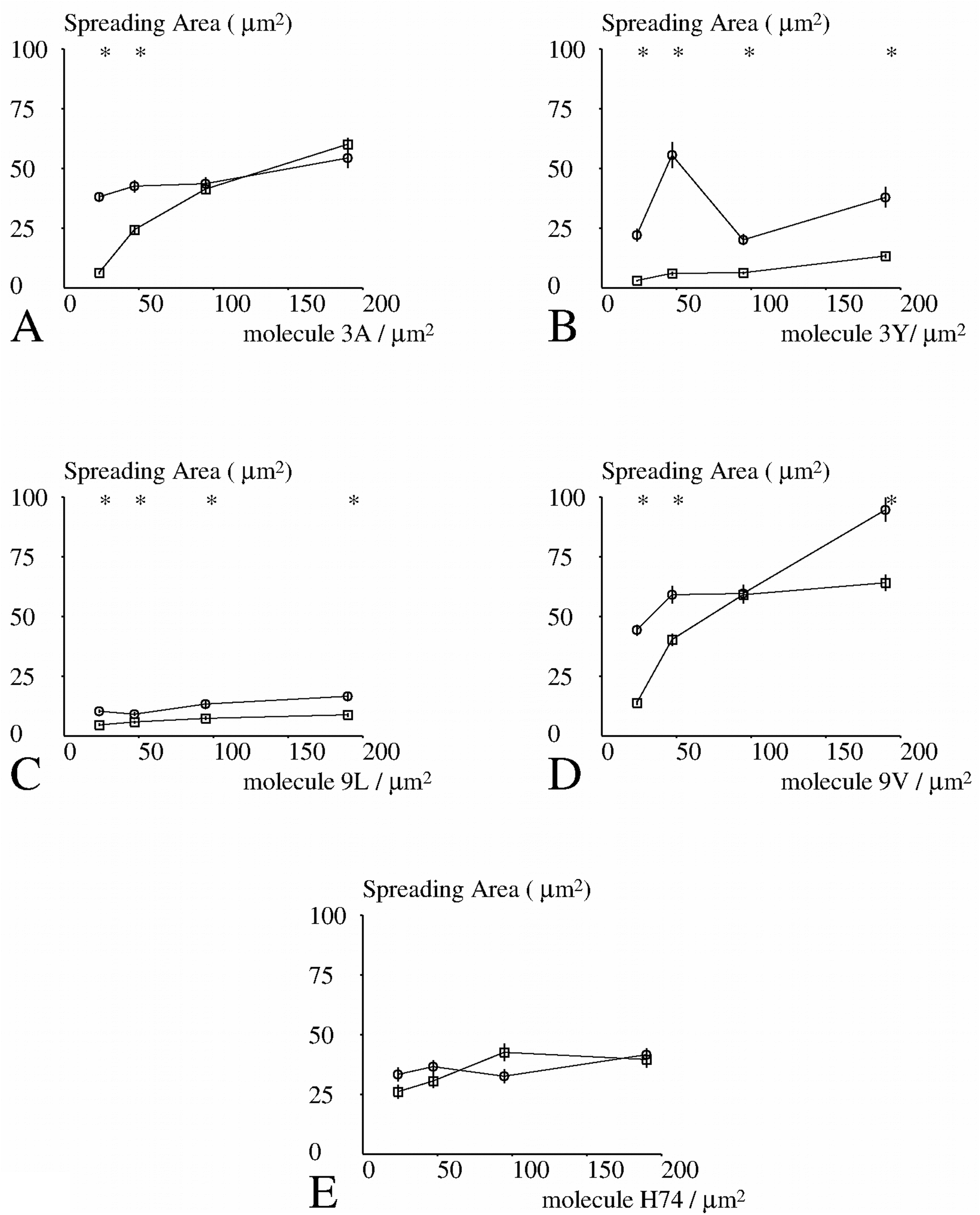
Dependence of spreading area on CD8 and activating peptide. CD8+ (open circles) or CD8− (open squares) Jurkat cells were deposited on surfaces bearing different amounts of peptide ligands for 1G4 and the contact area was determined 12-15 minutes after deposition. Results obtained after processing 13574 images are shown. Each point represents a mean of between 249 and 1096 values. Vertical bar length is twice the standard error for the mean. The significance of differences between CD8+ and CD8− cells was calculated with Student’s t test with Satterthwaite correction for determination of the number of degrees of freedom. Stars indicate a significance P< 0.001.

As shown of Figure 7, similar differences were obtained when the fraction of cells that could be triggered to spread was measured.

**Figure 7.**
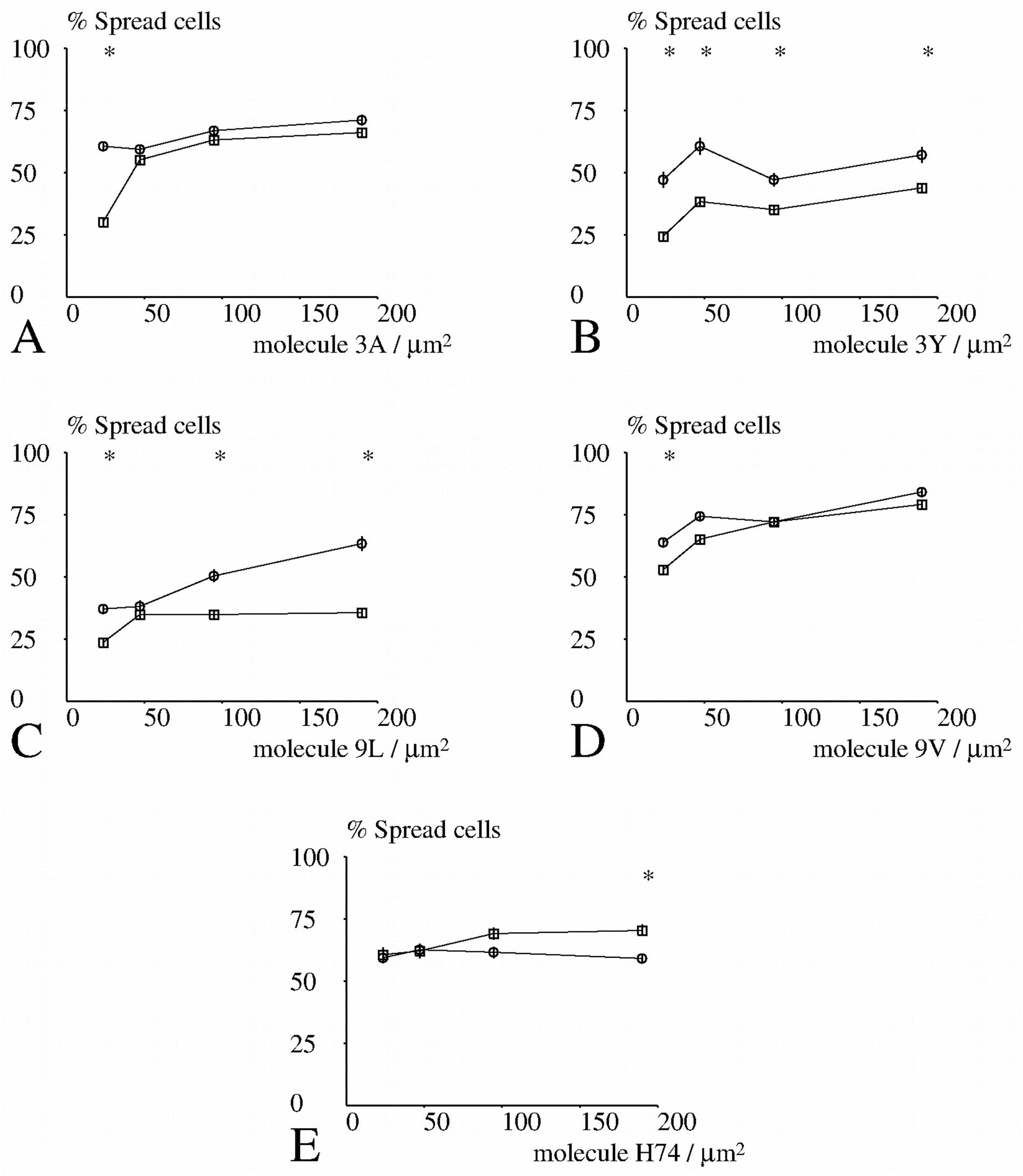
Dependence of the fraction of spreading cells on CD8 and activating peptide. CD8+ (open circles) or CD8− (open squares) cells were deposited on surfaces coated with different amounts of TCR ligands and the fraction of cells displaying significant spreading 12-15 minutes after deposition was measured. Results obtained after processing 13574 images are shown. Each point represents a mean of between 249 and 1096 values. Vertical bar length is twice the standard error for the mean. The significance of differences between CD8+ and CD8− cells was calculated with Student’s t test with Satterthwaite correction. Stars indicate a significance P< 0.001.

Thus, our results showed that CD8+ displayed higher sensitivity to spreading stimuli and more active spreading. However, the contact time required for spreading initiation was not lower.

Since CD8 was reported to influence binding as well as signaling events, it was important to determine whether the difference between the spreading behavior of CD8+ and CD8− cells was correlated to a difference between binding capacity. Since T cell activation was found to enhance binding capacity within seconds (*Pielak 2017*, *Hong 2018*), we studied the earliest steps of bond formation in order to try and observe activation-independent events.

### Cell adhesion under flow

#### CD8+ and CD8− cells display comparable binding capacity to pMHC coated surfaces

First, CD8+ and CD8− cells were driven along pMHC-coated surfaces and binding events were recorded. A total number of 3,936 arrests were detected after monitoring a total displacement length of 16.7 million pixels (i.e. 13.6 meters) corresponding to a mean arrest frequency of 0.3 per mm. In control experiments, the binding linear density (BLD) measured on uncoated surfaces was 0.027±0.02SE mm^−1^.

First, we used the sign test to perform a global comparison of the adhesive capacities of CD8+ and CD8− cells. In contrast with spreading experiments, no difference was found (P=0.65). Since the statistics of individual experimental conditions were not sufficient to detect limited differences between cells, we tentatively pooled the BLD obtained on the five peptide species. The BLD of CD8− and CD8+ cells were respectively 0.555 (± 0.022 SE) and 0.575 (± 0.022 SE) mm^−1^ on surfaces coated with 190 pMHC molecules/μm^2^ and respectively 0.133 (± 0.006 SE) and 0. 124 (± 0.004 SE) mm^−1^ on surfaces coated with 19 pMHC molecules/μm^2^.

##### CD8+ cells form bonds of similar duration as CD8− cells with pMHC-coated surfaces

Since the lifetime of binding events involving T cell receptors is considered as a key parameter of the activation process, it was important to know whether the spreading behaviour of CD8+ and CD8− cells was correlated to the lifetime of bonds formed with activating surfaces.

First, we performed a sign test to compare the fraction of attachments surviving 1s or 5 s after initial binding under different experimental conditions. No difference (P>0.6) was found between CD8− and CD8+ cells.

Secondly, since cell arrests were rare events (BLD lower than 1 mm^−1^), and arrest lifetime was of comparable order of magnitude as single bond lifetime (*Limozin 2019*), it seemed reasonable to assume that these arrests were mediated by single bonds. Therefore, we pooled the lifetime durations obtained with different pMHC surface densities to obtain sufficient statistics. As shown on Figure 8, survival curves yielded by CD8− and CD8+ cells were quite similar. Surprisingly, while no significant difference was found with 3Y, 9L, 9V and H74 (P>0.16 in all cases), the lifetime of bonds involving 3A was significantly lower with CD8+ than with CD8− cells (P<0.001).

**Figure 8.**
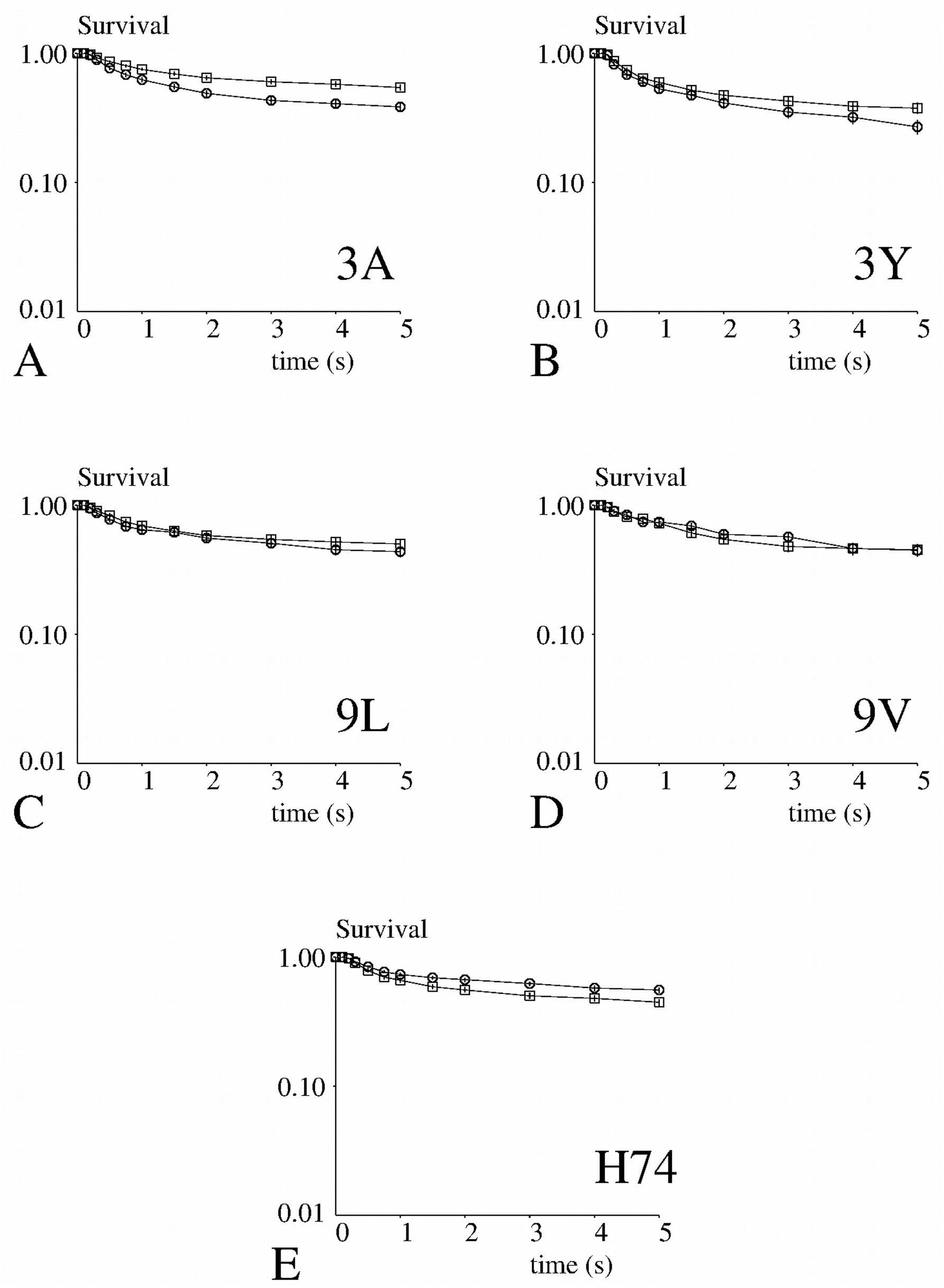
Dependence of bond lifetime on CD8 and activating peptide. CD8+ (open circles) or CD8− (open squares) were made to adhere to peptide-presenting surfaces in a laminar flow chamber and the duration of binding events was monitored for 5 seconds. The fraction attachments surviving at time t after formation was plotted versus time. Each curve represent between 153 and 444 binding events. Vertical bar length is twice the standard error as calculated (Methods)

Thus, the main conclusion of our experiments was that the higher spreading of CD8+ cells of activating surfaces as compared to CD8− cells could not be ascribed to a CD8-mediated increase of binding interactions

#### Measuring bonds beween TCRs and pMHCs in cell-free systems may provide a reasonable account of short term binding events involving whole cells

While standard methods of low shear hydrodynamics may provide an accurate account of the encounter conditions (contact duration and pulling force) of microsphere-bound and surface-bound ligands and receptors in a flow chamber, it is more difficult to estimate the precise properties of interactions involving living cells. Indeed, the pulling force is dependent on the nano- and micro-scale of interacting surfaces (*Goldman 1967*, *Pierres 1995*), and bond formation is dependent on the molecular environment of ligand and receptor molecules (*Pielak 2017*). It was therefore of high interest to compare our results with recent experiments performed on the same molecules in cell-free system (*Limozin 2019*). While cells were tested with a single wall shear rate, particles were subjected to five different shear rates, resulting in five different values of the average contact time between moving TCRs and fixed pMHCs, and five different values of the forces exerted by the flow on newly formed bonds. The correlation coefficient between adhesion of cells and particles were calculated for BLD (Figure 9A) and bond lifetime (Figure 9B). Clearly, the highest correlation for binding efficiency was found for an encounter duration of 1 ms (Figure 9A) and a disruptive force of 45 pN (Figure 9B). In addition to correlations, we compared the mean survival of bonds formed with different peptides. As shown on Figure 9C, the root mean square difference between survivals was on the order of 0.1. This has to be compared with survival of about 0.5.

**Figure 9.**
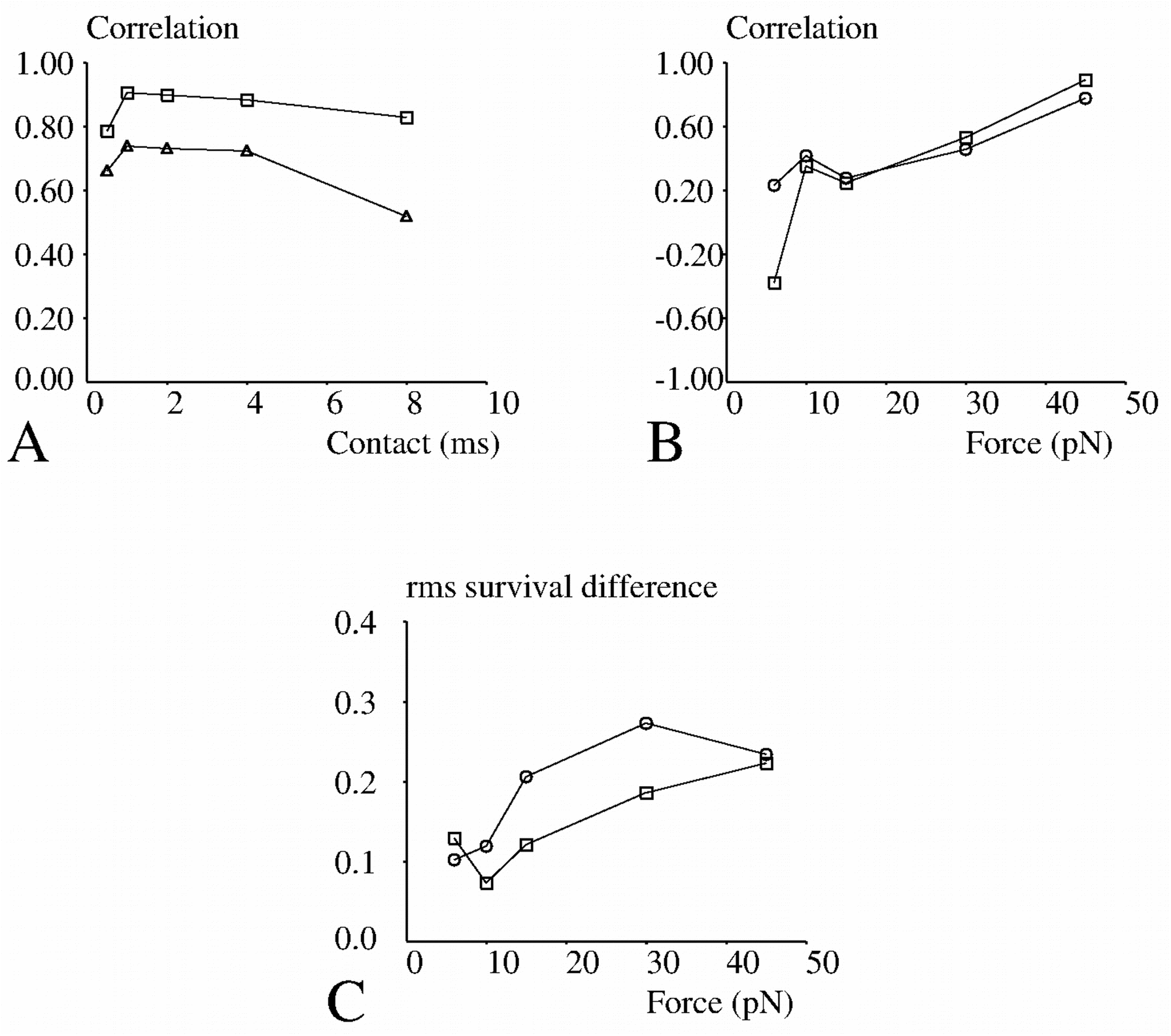
Comparison between the attachment behaviour of 1G4 bearing cells or microspheres under flow. A – The BLD under flow of cells on five different peptide species with a surface density of 190 molecules/μm^2^ (open squares) or 19 molecule/μm^2^ (open triangles) was determined, and the correlation coefficient was calculated between the obtained sets of five values and binding efficiencies measured on microspheres under various estimated durations of ligand-receptor contacts (as estimated from shear rate, *Limozin 2019*). The dependence of correlation coefficient on contact duration is shown. B – The mean survival of attachments formed with five peptide species were measured 1s (open squares) or 5 s (open circles) after binding. The correlation between these sets of five values and data obtained with microspheres (*Limozin 2019*) was calculated for various values of the pulling force applied to bonds. C – The root mean square of the differences between cell and microsphere bond survival at 1s (open squares) or 5 s (open circles) after binding were determined for various values of the pulling force applied to bonds after attachment.

## Discussion

The primary purpose of this work was to explore the role of CD8 in the detection and initial activation of T cells encountering cognate pMHCI ligand. Two parameters were studied : spreading of cells encountering activating surfaces and kinetics of bond formation and dissociation between TCRs and ligands. An additional output of our study might be to allow accurate comparison of the properties of cell surface receptors at the cell surface and in cell free conditions, which is important to assess the significance of model systems used to investigate cell functions (*Limozin 2019a*)

### Spreading experiments

Our strategy consisted of dropping CD8+ and CD8− T cells on surfaces coated with different amounts of TCR ligands with variable activation potency and measuring spreading kinetics. The rationale was that spreading was arguably the earliest cell-scale response triggered by TCR-ligand interaction; it was previously shown to be tightly correlated to a later proliferation program, which is considered as a robust reporter of T cell activation (*Cretel 2011*), and our setup allowed precise monitoring of cell behaviour immediately after contact with activating surfaces. Our experiments unambiguously showed that CD8+ cells displayed higher spreading activity than CD8− cells, and relative differences were more important when peptide surface densities were lowest (Figure 6).

It must be emphasized that formation of an extended contact is of high physiological relevance. Indeed, when two-photon microscopy was used to study the dynamics of antigen recognition in vivo, arrest and contact tightening were found to be the earliest consequences of antigen detection (*Cahalan 2008*). Two points deserve some discussion concerning the significance of our experiments.

First, the antigen sensitivity of T cells is known to depend on their state and environment : a cell migrating in a solid environment may react differently from a cell in suspension, due e.g. to shape changes (*Negulescu 1996*). The arrest of a moving cell encountering a cognate TCR ligand is certainly the earliest reporter of antigen detection and this arrest might be deemed a better reporter of antigen detection than spreading. However, the mechanisms of 2D and 3D migration are different (*Lammermann2008*), and while 3D migration is more representative of in vivo phenomena (*Cahalan 2008*), initial contacts and contact areas are difficult to measure accurately under 3D condition. Further, while 2D migration is easier to quantify (*Dustin 1997*) it may be less representative of physical conditions. Thus, observing the encounter between sedimenting cells and activating surfaces may provide a satisfactory trade-off between physiological relevance and liability to monitoring. This setup was indeed used in recent highly informative reports (*Lee 2017*, *Razvag 2019*).

Secondly, signaling events such as calcium spikes or recruitment and activation of kinases such as Zap70 were chosen as reporters of T cell activation. While much important information was obtained with these approaches (*Li 2004*), it must be recognized that interpretation is often ambiguous due to the formidable complexity of metabolic networks. Thus, while intracellular calcium is known to strongly influence many important processes (*Lee 2017*), intracellular spikes may occur spontaneously, and they may be linked to processes unrelated to *bona fide* activation. Also, the significance of calcium spikes observed seconds or minutes after TCR engagement may be different. Thus, monitoring mesoscale phenomena such as cell spreading may provide valuable information to understand the rationale of lymphocyte behaviour (*Gardel 2015, Malissen 2015*).

Indeed, the strong correlation between the kinetics and extent of cell spreading (Figure 3) supports the assumption that the spreading phenomenon is mainly driven by an internal cell program, in analogy with an hypothesis about the “stop and go” model of lymphocyte search for antigens (*Cahalan 2008*). The relative independence of the lag between cell-surface encounter and spreading initiation (Figure 4) would also be compatible with the hypothesis that that surface analysis might be driven by an internal program with relatively fixed duration.

Thus, it may be concluded that CD8+ cells displayed higher capacity than CD8− cells to trigger a spreading program after detecting cognate TCR ligands. It was important to determine within the framework of this accurately quantified model whether CD8 might act by enhancing TCR-pMHC interaction or by increasing T cell sensitivity to TCR engagement. We addressed this question by comparing the kinetics of initial TCR engagement and bond rupture in CD8+ and CD8-cells.

#### Adhesion experiments

We used a laminar flow chamber to study the frequency and lifetime of attachments between cells driven along surfaces coated with various surface densities (from 19 to 190 molecules/μm^2^) of the five pMHC species used for spreading experiments. No difference was found between the adhesive behavior of CD8+ and CD8− cells. These results make unlikely the validity in our model of two previously suggested hypotheses concerning the possible role of CD8 :

- First, our results did not support the hypothesis that CD8 might increase the kinetics of TCR engagement (*Wyer1999*, *Dutoit 2003*, *Gakamski 2005*).
- Second, since the survival of attachments was quantified up to 5 second duration, CD8 did not seem to increase bond stability during the first five seconds following initial attachment. This is consistent with a pioneer study made with atomic force microscopy, revealing that CD8 was not increasing adhesion efficiency for an activating peptide in a mouse system (*Puech 2011*). This may however seem at variance with recent reports that CD8 might increase the lifetime of TCR/pMHC interactions (*Kolawole 2018*, *Hong 2018*). However, while it seems well demonstrated that T cell activation may enhance the stability of the interaction between TCR born by CD8+ cells and cognate pMHCs (*O’Rourke 1990 & 1992*, *Hong 2018*, *Kolawole 2018*), it is not clear whether this phenomenon can occur during the first seconds following initial TCR engagement. Also, the effect of CD8 on TCR-pMHC interaction was reported to depend on tested TCR (*Zhong 2013*).

Thus, our experiments support the conclusion that CD8 enhanced the triggering of T cell functions after receptor engagement rather than facilitating TCR/pMHC interaction.

In addition, this report may bring important information concerning the influence of cell environment on the properties of membrane receptors.

#### Understanding the relationship between the binding properties of isolated and membrane-embedded receptors

It is now well recognized that the physical properties of biomolecule interactions are different when they occur between soluble molecules (i.e. under so-called 3D conditions) on between molecules bound to surfaces (2D conditions). However, we need more information to relate results from studies made on bonds formed between ligands and receptors bound to plain microparticles and associations between similar molecules embedded in cell membranes. Indeed, there is clear experimental support to the hypothesis that the cell environment may dynamically alter receptor function (*Pielak 2017*). It is therefore of high practical and theoretical interest to compare results of the present studies to recently reported properties on the interaction between microspheres bearing 1G4 TCR and surfaces exposing the same pMHC species as were used in the present study (*Limozin 2019*). In these studies, the binding efficiency and bond lifetime were studied under a range of shear rates allowing the encounter time between binding sites to vary between about 0.5 ms and 8 ms, and a force applied on bonds ranging between about 6 and 45 pN. The following points may be considered.

A first point is that the relative binding efficiencies and bond lifetimes of the five pMHC species were similar when TCRs were born by cells or microspheres (Figure 9 A&B). The correlation between both sets of results was highest when encounter time was close to 2 ms and distracting force was close to 45 pN, which is approximately consistent with the conditions of the present study (see Materials and Methods).

A second point is that the lifetime of attachments formed by cells was fairly comparable with values measured on microspheres (Figure 9C). This is a strong support to the assumption that the binding events recorded in our study were due to single molecular bonds.

A third point was that cell-surface binding was much less efficient than was found with microspheres : Indeed, while the binding linear density of microspheres to surfaces coated with about 1 μg/ml pMHC (yielding about 19 binding sites per μm^2^) was on the order of 100 mm^−1^ (*Limozin 2019)*, cells displayed more than one hundredfold lower BLD. Since TCRs were reported to be concentrated on the tip of T lymphocyte microvilli (*Jung 2016*), we think that the low efficiency of binding might be accounted by three processes : first, the macromolecular and particularly glycocalyx components might increase the nanoscale distance between TCRs and pMHCs as was demonstrated experimentally in other systems (*Sabri 2000*), secondly, the length and flexibility of the molecular structures bearing TCR and pMHC binding sites might be much lower in lymphocytes that on microspheres, resulting in a drastic reduction of the contact time between these sites during cell passage. Third, the overall effective density of receptors able to bind to the surface may be limiting in the case of cells due to the low surface ratio of microvilli tips versus average cell surface. On the contrary, typical conditions for microparticles are set to a saturating density of receptors on the particle surface (*Limozin 2019*). These findings raise two points that may deserve some discussion.

The first point is about the potential of the flow chamber and other devices to study the kinetics of unimolecular attachments. When tools such as atomic force microscopes (*Moy 1994*), biomembrane force probes (*Merkel 1999*), or optical traps (*Bartsch 2009*) are used to study single molecular bond, a major problem is demonstrating that observed binding events are actually representative of single molecular bonds. A fairly common argument consisted of adapting contact time in order than a low proportion of contacts might be conducive to bond formation, thus deducing from Poisson’s law that most binding events were mediated by single bonds. However, this reasoning is not fully rigorous, since it relies on the assumption that single bonds are actually detectable and that the “minimal detectable binding” is not, e.g. a bimolecular or a trimolecular attachment. We feel that a better proof of the single molecule assumption consists of performing sequential reduction of the surface density of binding sites and checking that i) BLD is proportional to the surface density, and ii) upon dilution, the lifetime distribution of binding events is unaltered. This check was extensively performed in our recent study of the interaction between 1G4 bearing microspheres and pMHC-coated surfaces (*Limozin 2019*). However, while cellular models have been studied for decades in our laboratory, performing the aforementioned check was not always feasible with these cellular systems for several reasons : i) since cells are usually more heterogeneous than microspheres, and different cells may have different binding abilities, ligand dilution may decrease the BLD of only a fraction of tested cells. It may be difficult to obtain a linear relationship between ligand surface density and binding efficiency. ii) if cell binding ability is too low, it may not be feasible to decrease the surface concentration of ligands sufficiently to achieve a linear proportion between BLD and surface density, since nonspecific binding events may become too frequent. It must also be pointed out that non specific interactions may prevent the use of too low shear rates. This is certainly the reason why flow chambers met with prominent success when they were first applied to selectin-mediated interactions that displayed a unique capacity to form in presence of high shear rates as may be found in flowing blood (*Lawrence 1991*). However, in the present case, the strong reduction of BLD for cell experiments as compared with microparticle experiments (where single bond was fully assessed) is an additional indication that single bonds are also measured with cells.

The second point is about the significance of so-called 2D affinity. While the conventional affinities of biomolecule interactions as measured in bulk liquid phase are fairly reproducible and are not too dependent on experimental conditions, interactions between surface bound molecules are much more sensitive to poorly controlled and highly variable parameters such as distance between surfaces, nanoscale surface fluctuations, and presence of repeller macromolecules. Indeed, the above comparison suggests that 2D affinity may display hundredfold variations due to molecular environment. Thus, the concept of 2D affinity as an intrinsic property of a biomolecule couple may be quite misleading. In this context, taking into account the distance between membranes, helps reconciliating observations between particle and membrane-born receptors, as recently illustrated for CD16-mediated NK cell spreading (*Gonzalez 2019*).

A last point is about the link between our results and our understanding of the strategy followed by lymphocytes to scan living organisms in order to detect as rapidly as possible the presence of foreign structures. It has long been recognized that this requires outstanding performances that might seem incompatible with limitations set by physical laws (*Malissen15*) : indeed, since T lymphocytes specific for a given antigen are highly outnumbered by antigen presenting cells in lymph nodes, it is not surprising that contact need to be quite short. Average duration was estimated at 3.7 minutes with two-photon microscopy (*Miller 2004*). Interestingly, this is the same order of magnitude as the time required for all pMHCs exposed by an antigen-presenting cell to pass through a small contact area such as the tip of a cell protrusion (*Bongrand 1998*). However, this timescale is much longer than the transit time of a pMHC through a small contact area (this is on the order of 0.1s estimating the diffusion rate as 100 μm^2^/s and the size of the contact area as 1 μm^2^). This is consistent with recent report showing that lymphocytes may scan a neighbouring surfaces with highly dynamic microvilli generating molecular contacts of a few second duration (*Brodovitch 2013*, *Cai 2017*). After a contact of a few minutes, the cell may take the decision to tighten the contact for more accurate analysis or depart to examine another cell.

The hypothesis suggested by our experiments would be that during the initial minute-scale phase cells might sum the properties of individual molecular contacts and take a decision that might account for the longest contact duration or the total contact duration, depending on stochastic variations of individual interactions (*Lin 2019*). That the time of spreading initiation was fairly independent of pMHCs is consistent with the view that either a 2-minute lag is fully determined by an internal cell program or alternatively that the initial analysis is too crude to discriminate between pMHCs of slightly different activation potency. The dependence of the spreading rate on the peptide suggests that a two minute lag is sufficient for the cell to acquire quantitative information on the surface it has been analyzing.

In conclusion, we suggest that a quantitative analysis of mesoscale functioning is required to unravel the formidable complexity of signaling events involved in T cell activation. This complexity is well illustrated by the following two examples : in a study of the effect of myosin II on the maturation of focal adhesions, 905 protein species were found to be involved and 459 of these changed in abundance when myosin was inhibited (*Kuo 2011*). In a recent proteomic study of T lymphocyte activation, 112 high-confidence molecular interaction were found to be involved (*Roncagalli 2014*). In accordance with recent reports (*Gardel 2015, Malissen 2015*), we suggest that a quantitative study of the dependence of T Cell Antigen Recognition on T Cell Receptor-Peptide MHC confinement time to understand mesocale cell functions is a prerequisite to an understanding of the data provided by high-throuput studies.

## Acknowledgment

The excellent technical assistance of Dominique Touchard is gratefully acknowledged. The authors declare no conflict of interest.

